# Symbiotic and non-symbiotic members of the genus *Ensifer* (syn. *Sinorhizobium*) are separated into two clades based on comparative genomics and high-throughput phenotyping

**DOI:** 10.1101/2020.04.23.055285

**Authors:** Camilla Fagorzi, Alexandru Ilie, Francesca Decorosi, Lisa Cangioli, Carlo Viti, Alessio Mengoni, George C diCenzo

**Affiliations:** Department of Biology, University of Florence, Sesto Fiorentino, Italy; Genexpress Laboratory, Department of Agriculture, Food, Environment and Forestry, University of Florence, Sesto Fiorentino, Italy; Department of Biology, Queen’s University, Kingston, Ontario, Canada

**Keywords:** Mutualism, evolutionary biology, phenomics, comparative genomics, rhizobia, Proteobacteria

## Abstract

Rhizobium – legume symbioses serve as a paradigmatic example for the study of mutualism evolution. The genus *Ensifer* (syn. *Sinorhizobium*) contains diverse plant-associated bacteria, a subset of which can fix nitrogen in symbiosis with legumes. To gain insights into the evolution of symbiotic nitrogen fixation (SNF), and inter-kingdom mutualisms more generally, we performed extensive phenotypic, genomic, and phylogenetic analyses of the genus *Ensifer*. The data suggest that SNF emerged several times within the genus *Ensifer*, likely through independent horizontal gene transfer events. Yet, the majority (105 of 106) of the *Ensifer* strains with the *nodABC* and *nifHDK* nodulation and nitrogen fixation genes were found within a single, monophyletic clade. Comparative genomics highlighted several differences between the “symbiotic” and “non-symbiotic” clades, including divergences in their pangenome content. Additionally, strains of the symbiotic clade carried 325 fewer genes, on average, and appeared to have fewer rRNA operons than strains of the non-symbiotic clade. Characterizing a subset of ten *Ensifer* strains identified several phenotypic differences between the clades. Strains of the non-symbiotic clade could catabolize 25% more carbon sources, on average, than strains of the symbiotic clade, and they were better able to grow in LB medium and tolerate alkaline conditions. On the other hand, strains of the symbiotic clade were better able to tolerate heat stress and acidic conditions. We suggest that these data support the division of the genus *Ensifer* into two main subgroups, as well as the hypothesis that pre-existing genetic features are required to facilitate the evolution of SNF in bacteria.

## INTRODUCTION

Symbioses are pervasive phenomena present in all Eukaryotic forms of life (López-García et al. 2017). These includes the evolution of organelles, obligate symbioses, and facultative symbiotic interactions (Douglas 2014), with symbiotic nitrogen fixation (SNF) being a paradigmatic example of the latter (Masson-Boivin & Sachs 2018). SNF (the conversion of N_2_ to NH_3_) is performed by a polyphyletic group of bacteria from the Alphaproteobacteria and Betaproteobacteria (whose nitrogen-fixing members are collectively called rhizobia) and members of the genus *Frankia* (Wang & Young 2019; Masson-Boivin et al. 2009) while intracellularly housed within specialized organs (nodules) of specific plants in the family *Fabaceae* and the genus *Parasponia*, as well as the actinorhizal plants (Werner et al. 2014; Velzen et al. 2018; Griesmann et al. 2018). The advantages and evolutionary constraints to SNF have long been investigated in the conceptual framework of mutualistic interactions and the exchange of goods (see for instance (Sørensen et al. 2019; Werner et al. 2015; Heath & Tiffin 2007)), and quantitative estimations with metabolic reconstructions have also been performed (Pfau et al. 2018; diCenzo, Tesi, et al. 2019).

The establishment of a symbiotic nitrogen-fixing interaction requires that the bacterium encode several diverse molecular functions, including those related to signalling and metabolic exchange with the host plant, nitrogenase and nitrogenase-related functions, and escaping or resisting the plant immune system (Oldroyd et al. 2011; Haag et al. 2013; Poole et al. 2018). In general, the core SNF genes (*nod, nif, fix* genes, among others) are located within mobile genetic elements that include symbiotic islands and symbiotic (mega)-plasmids (Tian & Young 2019; Checcucci et al. 2019; Geddes et al. 2020), facilitating their spread through horizontal gene transfer (HGT) (Sullivan et al. 1995; Barcellos et al. 2007; Pérez Carrascal et al. 2016). Emphasizing the role of HGT in the evolution of rhizobia, rhizobia are dispersed across seven families of the Alphaproteobacteria and one family of the Betaproteobacteria, and most genera with rhizobia also contain non-rhizobia (Garrido-Oter et al. 2018; Wang 2019).

An interesting area of investigation is whether the evolution of mutualistic symbioses, such as SNF, depends on metabolic/genetic requirements (“facilitators”, as in (Gerhart & Kirschner 2007)) aside from the strict symbiotic genes (Zhao et al. 2017; Long 2001; Sanjuán 2016). In other words, i) is the acquisition of symbiotic genes present in genomic islands or plasmids sufficient to become a symbiont or, ii) are metabolic pre-requirements or adaptation successive to HGT required? A comparative genomics study of 1,314 Rhizobiales genomes identified no functional difference between rhizobia and non-rhizobia based on Kyoto Encyclopedia of Genes and Genomes (KEGG) annotations (Garrido-Oter et al. 2018), suggestive of an absence of obvious facilitators. In contrast, experimental studies are generally consistent with an important role of non-symbiotic genes in establishing or optimizing rhizobium – legume symbioses. Several studies have shown that effective symbionts are not produced following the transfer of symbiotic plasmids from rhizobia of the genera *Rhizobium* or *Ensifer* (syn. *Sinorhizobium*) to closely related non-rhizobia from the genera *Agrobacterium* or *Ensifer* (see for instance (Hooykaas et al. 1982; Finan et al. 1986; Rogel et al. 2001); reviewed in (diCenzo, Zamani, et al. 2019)). Similarly, the same symbiotic island is associated with vastly different symbiotic phenotypes depending on the *Mesorhizobium* genotype (Nandasena et al. 2007; Haskett et al. 2016). Further supporting the need for additional adaptations to support SNF, symbiosis plasmid transfer coupled to experimental evolution can lead to the gain of more advanced symbiotic phenotypes (Doin de Moura et al. 2020).

The genus *Ensifer* provides an ideal model to further explore the differentiation, or lack thereof, of symbiotic bacteria from non-symbionts. This genus comprises rhizobia such as *E. meliloti* and *E. fredii*, as well as non-rhizobia like *E. morelense* and *E. adhaerens*, and many members have been extensively studied producing an abundant set of experimental and genomic data (for a recent review, see (diCenzo, Zamani, et al. 2019)). The genus *Ensifer*, as currently defined, resulted from the combination of the genera *Sinorhizobium* and *Ensifer* based on similarities in the 16S rRNA and *recA* sequences of the type strains and the priority of the name *Ensifer* (Young 2003; Willems et al.). Multilocus sequence analysis supported the amalgamation of these genera (Martens et al. 2007), although it was subsequently noted that *E. adhaerens* (the type strain) is an outgroup of this taxon based on whole genome phylogenomics (Ormeño-Orrillo et al. 2015). A more recent taxonomy approach based on genome phylogeny suggests that the genus *Ensifer* should again be split, with the initial type strains of *Ensifer* and *Sinorhizobium* belonging to separate genera (Parks et al. 2018).

In this paper we report an extensive comparative genomic and phenotypic characterization of legume symbionts and non-symbionts of the genus *Ensifer*. We identified that SNF likely evolved multiple times through independent HGT events; even so, most symbionts were found in a single clade, consistent with a requirement for pre-existing genetic features to facilitate the evolution of SNF. Moreover, the symbiotic and non-symbiotic clades differed in their pangenome composition as well as their substrate utilization and resistance phenotypes as measure by the Phenotype MicroArray™platform. We suggest that the data support the division of the genus *Ensifer* into two subgroups, corresponding to the genera *Ensifer* and *Sinorhizobium* of the Genome Taxonomy Database (Parks et al. 2018).

## MATERIALS AND METHODS

### Genome Sequencing, Assembly, and Annotation

Prior to short-read sequencing, all strains were grown to stationary phase at 30°C in TY medium (5 g L^-1^ tryptone, 3 g L^−1^ yeast extract, and 0.4 g L^-1^ CaCl_2_). Total genomic DNA was isolated using a standard cetyltrimethylammonium bromide (CTAB) method (Perrin et al. 2015). Short-read sequencing was performed at IGATech (Udine, Italy) using an Illumina HiSeq2500 instrument with 125 bp paired-end reads. Two independent sequencing runs were performed for *E. morelense* Lc04 and *E. psoraleae* CCBAU 65732, whereas *E. morelense* Lc18 and *E. sesbaniae* CCBAU 65729 were sequenced once. For the long-read sequencing, *E. sesbaniae* was grown to mid-exponential phase at 30°C in MM9 minimal medium (MOPS buffer [40 mM MOPS, 20 mM KOH], 19.2 mM NH_4_Cl, 85.6 mM NaCl, 2 mM KH_2_PO_4_, 1 mM MgSO_4_, 0.25 mM CaCl_2_, 1 µg ml^-1^ biotin, 42 nM CoCl_2_, 38 µM FeCl_3_, 10 µM thiamine-HCl, and 10 mM sucrose). Total genomic DNA was isolated as described elsewhere (Cowie et al. 2006). Long-read sequencing was performed in-house with a Pacific Biosciences Sequel instrument.

Reads were assembled into scaffolds using SPAdes 3.9.0 (Bankevich et al. 2012; Vasilinetc et al. 2015); in the case of *E. sesbaniae*, long reads were corrected and trimmed using Canu 1.7.1 (Koren et al. 2017) prior to assembly. Scaffolds returned by SPAdes were parsed to remove those with less than 20x coverage or with a length below 200 nucleotides. Using FastANI (Jain et al. 2018), one-way average nucleotide identity (ANI) values of each assembly were calculated against 887 alpha-proteobacterial genomes available through the National Center for Biotechnological Information (NCBI) with an assembly level of complete or chromosome. Based on the FastANI output, each draft genome assembly was further scaffolded using MeDuSa (Bosi et al. 2015) and the reference genomes listed in Supplementary Table S1. For most assemblies, scaffolds under 1 kb in length were discarded. The exception was for *S. sesbaniae*, for which case scaffolds under 10 kb were discarded. Genome assemblies were annotated using Prokka 1.12-beta (Seemann 2014), annotating coding regions with Prodigal (Hyatt et al. 2010), tRNA with Aragon (Laslett & Canbäck 2004), rRNA with Barrnap (github.com/tseemann/barrnap), and ncRNA with Infernal (Kolbe & Eddy 2011) and Rfam (Kalvari et al. 2018).

### Species Phylogenetic Analyses

All *Ensifer* (and *Sinorhizobium*) genomes were downloaded from the NCBI Genome Database regardless of assembly level. Strains that either i) lacked a RefSeq assembly, ii) had genome sizes < 1 Mb, or iii) appeared to not belong to the *Ensifer* clade based on preliminary phylogenetic analyses were discarded, leaving a final set of 157 strains (Supplementary Dataset S1). Eight complete *Rhizobium etli* genomes (Supplementary Dataset S2) were downloaded to serve as an outgroup. Genomes were reannotated with prokka to ensure consistent annotation.

To construct an unrooted, whole-genome phylogeny, the pangenome of the 157 *Ensifer* strains was calculated using Roary 3.11.3 (Page et al. 2015) with a percent identify threshold of 70%. The nucleotide sequences of the 1,049 core genes were individually aligned with PRANK (Löytynoja 2014). Alignments were concatenated and used to construct a maximum likelihood phylogeny (the bootstrap best tree following 50 bootstrap replicates) using RAxML 8.2.9 (Stamatakis 2014) with the GTRCAT model. All phylogenies prepared in this study were visualized with the online iTOL webserver (Letunic & Bork 2016).

To construct a rooted phylogeny, AMPHORA2 (Wu & Scott 2012) was used to identify 30 highly-conserved, single-copy proteins (Frr, InfC, NusA, Pgk, PyrG, RplA, RplB, RplC, RplD, RplE, RplF, RplK, RplL, RplM, RplN, RplP, RplS, RplT, RpmA, RpoB, RpsB, RpsC, RpsE, RpsI, RpsJ, RpsK, RpsM, RpsS, SmpB, Tsf) in each *Ensifer* and *Rhizobium* proteome, based on hidden Markov models (HMMs) and HMMER 3.1b2 (Eddy 2009). Orthologous groups were aligned using MAFFT 7.310 (Katoh & Standley 2013) with the localpair option, following which the alignments were trimmed using TRIMAL 1.2rev59 (Capella-Gutiérrez et al. 2009) with the automated1 option. Alignments were concatenated and used to construct a maximum likelihood phylogeny (the bootstrap best tree following 300 bootstrap replicates) using RAxML with the PROTGAMMALG model.

### ANI and AAI Calculations

Pairwise ANI values were calculated for all *Ensifer* strains using FastANI (Jain et al. 2018) with default parameters; a value of 78% was used in cases where no value was returned by FastANI. Pairwise average amino acid identity (AAI) values were calculated with the compareM workflow (github.com/dparks1134/CompareM). Results were visualized and clustered using the heatmap.2 function of the gplots package in R (Warnes et al. 2016), with average linkage and Pearson correlation distances.

### Pangenome Calculation

All proteins of the reannotated *Ensifer* strains were clustered into orthologous groups using CD-HIT 4.6 (Li & Godzik 2006) with a percent identity threshold of 70% and an alignment length of 80% of the longer protein. The output was used to determine core and accessory genomes using a prevalence threshold of 90% as many of the genomes were draft genomes. Gene accumulation curves were produced using the specaccum function of the vegan package of R (Oksanen et al. 2018), with the random method and 500 permutations. Principal component analysis (PCA) was performed with the prcomp function of R, and was visualized with the autoplot function the ggplot2 package (Wickham 2016).

### Identification and Phylogenetic Analysis of Common Nod, Nif, and Rep Proteins

The proteomes were collected for the 157 *Ensifer* strains, as well as all strains from the genera *Rhizobium, Neorhizobium, Agrobacterium, Mesorhizobium*, and *Ochrobactrum* with an assembly status of Complete or Chromosome (Supplementary Dataset S3). Additionally, the seed alignments for the HMMs of the nodulation proteins NodA (TIGR04245), NodB (TIGR04243), and NodC (TIGR04242), the nitrogenase proteins NifH (TIGR01287), NifD (TIGR01282), NifK (TIGR01286), and the replicon partitioning proteins RepA (TIGR03453), and RepB (TIGR03454) were downloaded from TIGRFAM (Haft et al. 2013). Seed alignments were converted into HMMs with the HMMBUILD function of HMMER 3.1b2 (Eddy 2009). Each HMM was searched against the complete set of proteins from all 157 reannotated *Sinorhizobium* and *Ensifer* strains using the HMMSEARCH function of HMMER. The amino acid sequences for each hit (regardless of e-value) were collected. Each set of sequences was searched against a HMM database containing all 21,200 HMMs from the Pfam (Finn et al. 2016) and TIGRFAM databases using the HMMSCAN function of HMMER, and the top scoring HMM hit for each query protein was identified. Proteins were annotated as NodA, NodB, NodC, NifH, NifD, NifK, RepA, or RepB according to Supplementary Table S2.

The NodA, NodB, and NodC proteins were associated to operons based on identifying proteins that are encoded by adjacent genes in their respective genomes; proteins not found in a predicted operon encoding all three proteins were discarded. Each set of orthologs were aligned using MAFFT with the localpair option, and alignments trimmed using TRIMAL and the automated1 algorithm. Alignments were concatenated so as to combine alignments for proteins belonging to the same operon, producing a NodABC alignment. The same procedure was followed to produce NifHDK and RepAB alignments. Maximum likelihood phylogenies were built on the basis of each combined alignment using RAxML with the PROTGAMMALG model. The final phylogenies are the bootstrap best trees following 300, 400, and 350 bootstrap replicates for the NodABC, NifHDK, and RepAB phylogenies, respectively.

### Plant Assays

*Phaseolus vulgaris* (var. TopCrop, Mangani Sementi, Italy) seeds were surface sterilized in 2.5% HgCl_2_ solution for two minutes and washed five times with sterile water. Seeds were germinated in the dark at 23°C, following which seedlings were placed in sterile vermiculite:perlite (1:1) polypropylene jars and grown at 23°C with a 12 hour photoperiod (100 µE m^-2^ s^-1^). One-week old plantlets were inoculated with 100 µL of the appropriate rhizobium strain (suspended in 0.9% NaCl at an OD_600_ of 1); five plants were inoculated per strain and then grown for four weeks at 23°C with a 12 hour photoperiod (100 µE m^-2^ s^-1^). Nodules were collected and surface sterilized as described elsewhere (Checcucci et al. 2016), crushed in sterile 0.9% NaCl solution, and serial dilutions were plated on TY agar plates and incubated at 30°C for two days. Amplified Ribosomal DNA Restriction Analysis (ARDRA) was performed using lysates from single colonies. Briefly, the 16S rRNA gene was amplified with primers 27f and 1495r (Barzanti et al. 2007), and the product individually digested with 20 units of either *Msp*I or *Hha*I for 3 hours at 37°C.

### Phenotype MicroArray™

Phenotype MicroArray™experiments using Biolog plates PM1 and PM2A (carbon sources), PM9 (osmolytes), and PM10 (pH) were performed as described previously (Biondi et al. 2009). Data were collected over 96 hours with an OmniLog™instrument. Data analysis was performed with DuctApe (Galardini et al. 2014). Activity index (AV) values were calculated following subtraction of the blank well from the experimental wells. Growth with each compound was evaluated with AV values from 0 (no growth) to 9 (maximal growth), following an elbow test calculation.

### Biofilm Assays

Overnight cultures of strains grown in TY and LB (10 g L^-1^ tryptone, 5 g L^-1^ yeast extract, 5 g L^-1^ NaCl) media were diluted to an OD_600_ of 0.02 in fresh media, and six replicates of 100 µL aliquots were transferred to a 96-well microplate. Plates were incubated at 30°C for 24 hours, after which the OD_600_ was measured with a Tecan Infinite 200 PRO (Switzerland). Each well was then stained with 30 µL of a filtered 0.1% (w/v) crystal violet solution for 10 minutes. Medium containing the planktonic cells was gently removed, following which the wells were rinsed three times with 200 μL of phosphate-buffered saline (PBS; 0.1 M, pH 7.4) and allowed to dry for 15 min. One hundred µL of 95% (v/v) ethanol was added to each well and then incubated for 15 minutes at room temperature. The OD_540_ of each well was measured (Rinaudi & González 2009), and biofilm production reported as the ratio of the OD_540_/OD_600_ ratio.

### Indole-3-Acetic Acid (IAA) Production Assays

Two hundred µL of overnight bacterial cultures were transferred 5 mL of a 1:10 dilution of TSB medium (Bio-Rad, USA) supplemented with 1 mg mL^-1^ L-tryptophan and incubated at 30°C overnight (Gordon & Weber 1951). One hundred µL aliquots of each culture were transferred to a 96-well microplate, in triplicate, and OD_600_ values determined with a Tecan Infinite 200 PRO. Aliquots of each culture were centrifuged for three minutes at 10,000 *g*, and 50 µL of the supernatant was mixed with 50 µL of Salkowsky reagent (35% v/v perchloric acid, 10 mM FeCl_3_). The OD_530_ of each well was measured following 30 minutes of incubation at room temperature, and IAA production reported as the ratio of the OD_530_/OD_600_ ratio. *Escherichia coli* DH5α was used as a positive control, and uninoculated medium as the negative control.

### Growth Curves

Overnight cultures of each strain were grown in the same medium to be used for the growth curve. For minimal media, either 0.2% (w/v) of glucose or succinate was added as the carbon source. Cultures were diluted to an OD_600_ of 0.05 in the same media, and triplicate 150 µL aliquots were added to a 96-well microplate. Microplates were incubated without shaking at 30°C or 37°C in a Tecan Infinite 200 PRO, with OD_600_ readings taken every hour for 48 hours. Growth rates were evaluated over two-hour windows during the exponential growth phase.

To evaluate bacterial growth when provided root exudates as a nitrogen source, root exudates were produced from *Medicago sativa* cv. Maraviglia as described elsewhere (Checcucci et al. 2017). Single bacterial colonies from TY plates were resuspended in a 0.9% NaCl solution to an OD_600_ of 0.5. Then, each well of a 96-well microplate was inoculated with 75 µL of nitrogen-free M9 with 0.2% (w/v) succinate as a carbon source, 20 µL of root exudate as a nitrogen-source, and 5 µL of culture. Triplicates were performed for each strain. Microplates were incubated without shaking at 30°C in a Tecan Infinite 200 PRO, with OD_600_ readings taken every hour for 48 hours. Growth rates were determined as described above.

### Data Availability

Genome assemblies were deposited to NCBI under the BioProject accession PRJNA622509. Scripts to repeat the computational analyses reported in this study are available at: github.com/diCenzo-GC/Ensifer_phylogenomics.

## RESULTS

### Genome sequencing of four *Rhizobiaceae* strains

Draft genomes of *E. morelense* Lc04, *E. morelense* Lc18, *E. sesbaniae* CCBAU 65729, and *E. psoraleae* CCBAU 65732 (Wang et al. 1999, 2013) were generated to increase the species diversity available for our analyses. Summary statistics of the assemblies are provided in Supplementary Table S3. The genome sequences confirmed the presence of nodulation and nitrogen-fixing genes in *E. morelense* Lc18, *E. sesbaniae* CCBAU 65729, and *E. psoraleae* CCBAU 65732, while these genes appeared absent in the *E. morelense* Lc04 assembly. ANI values confirmed that strains Lc04, CCBAU 65729, and CCBAU 65732 belonged to the genus *Ensifer*. However, strain Lc18 was most similar to genomes from the genera *Rhizobium* and *Agrobacterium*, consistent with an earlier 16S rRNA gene restriction fragment length polymorphism analysis (34). Thus, we propose renaming *E. morelense* Lc18 to *Rhizobium* sp. Lc18. As this strain does not belong to the genus *Ensifer*, it was excluded from further analyses.

### Symbiotic and non-symbiotic *Ensifer* strains segregate phylogenetically

An unrooted, whole-genome phylogeny of 157 *Ensifer* strains was prepared to evaluate the phylogenetic relationships between the symbiotic and non-symbiotic strains (Figure 1). A rooted phylogeny based on a multi-locus sequence analysis was also prepared (Supplementary Figure S1). Each of the 157 strains were annotated as symbiotic or non-symbiotic based on the presence of the common *nodABC* nodulation genes and the *nifHDK* nitrogenase genes. Consistent with previous work (Garrido-Oter et al. 2018), both phylogenies revealed a clear division of the symbiotic and non-symbiotic strains into two well-defined clades. However, a few exceptions were noted. *E. sesbaniae*, a known symbiont (Wang et al. 2013) whose ability to effectively nodulate *Phaseolus vulgaris* was confirmed (Supplementary Figure S2), was found within the non-symbiotic clade. Similarly, at least one of the six symbiotic proteins were not detected in eight strains of the symbiotic group, although we cannot rule out that these are false negatives in our annotation pipeline. ANI and AAI calculations suggested the presence of 12 and 20 genospecies within the non-symbiotic and symbiotic groups, respectively (Figure 1, Supplementary Figures S3, S4), confirming that the non-symbiotic clade was not an artefact of low species diversity. Thus, we conclude that the genus *Ensifer* consists of two well-defined clusters, each consisting predominately of either symbiotic or non-symbiotic strains.

**Figure 1.**
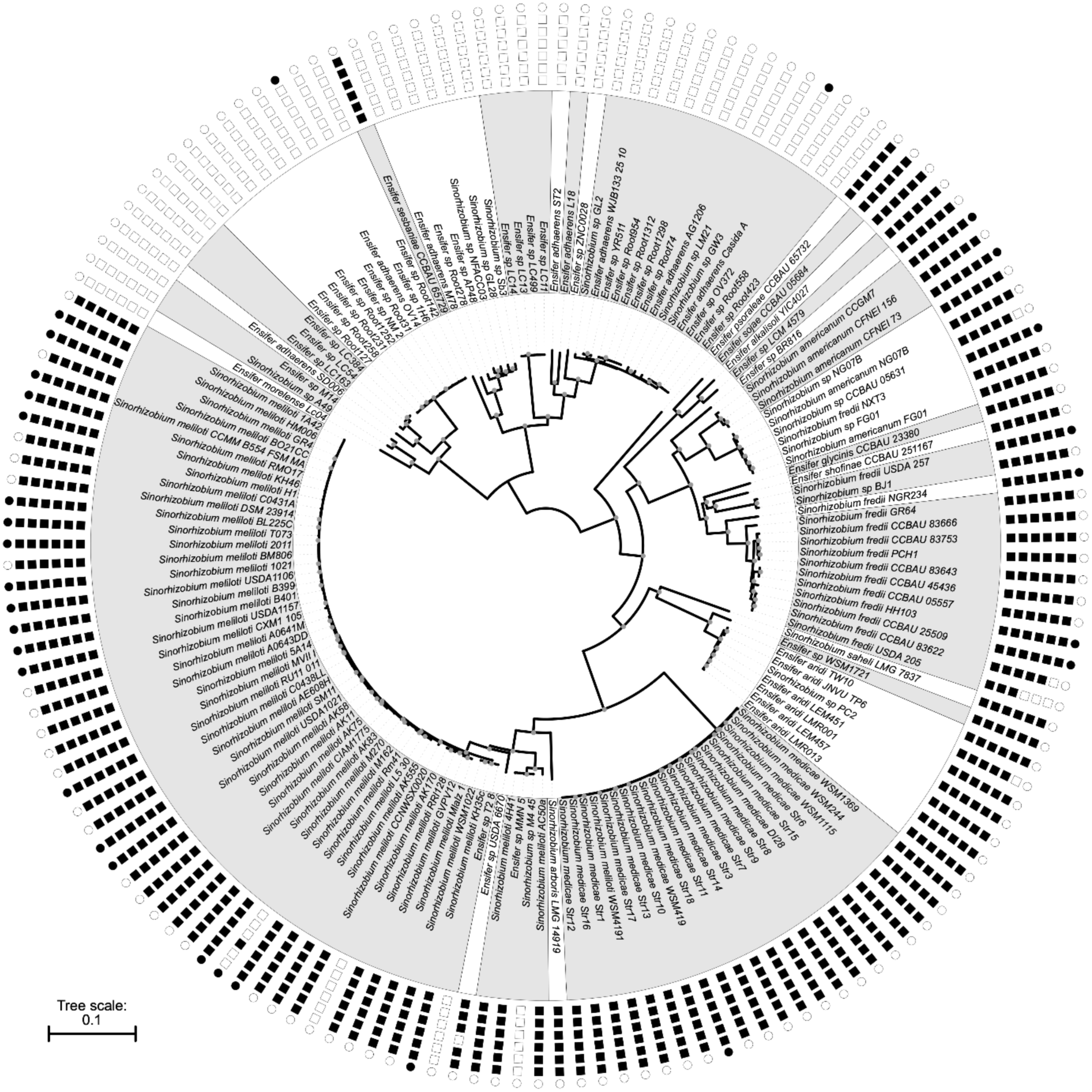
Unrooted phylogeny of the genus *Ensifer*. A maximum likelihood phylogeny of 157 strains was prepared from a concatenated alignment of 1,049 core genes. Nodes with a bootstrap value of 100 are indicated with the gray dots. The scale represents the mean number of nucleotide substitutions per site. The white and grey shading is used to group strains into genospecies on the basis of ANI and AAI results (Supplementary Figures S3, S4). From outside to inside, the outer rings represent the genome assembly level (black – finished, white – draft), and the presence (black) or absence (white) of NodA, NodB, NodC, NifH, NifD, and NifK. Strains are named as recorded in NCBI at the time of collection.

### SNF arose multiple times within the genus *Ensifer*

A possible explanation for the phylogenetic segregation of SNF within the genus *Ensifer* was that the symbiotic genes were gained once through a single HGT event. To test this hypothesis, the phylogenetic relationships of the NodABC and NifHDK proteins of the order Rhizobiales were examined (Figures 2A, 2B). SNF genes are situated on megaplasmids in the genus *Ensifer*; thus, a phylogeny of RepAB partitioning proteins of the order Rhizobiales was prepared as a proxy of the evolutionary relationships among the symbiotic megaplasmids (Figure 2C). All three phylogenies were inconsistent with a single origin of SNF within the genus *Ensifer* as the *Ensifer* strains were predominantly split into three clades: i) *E. meliloti* and *E. medicae*, ii) *E. fredii* and related strains, and iii) *E. americanum* and related strains. As the same clades are observed in the species tree (Figure 2D), this observation suggests SNF was independently acquired in each clade. The relationships between the SNF genes of the remaining *Ensifer* species (e.g., *E. aridi* and *E. psoraleae*) was not clear; however, the most parsimonious solution is that there were three additional acquisitions of SNF (Figure 2D). Taken together, these data suggest that SNF arose several independent times in the genus *Ensifer*, but preferentially within one monophyletic clade.

**Figure 2.**
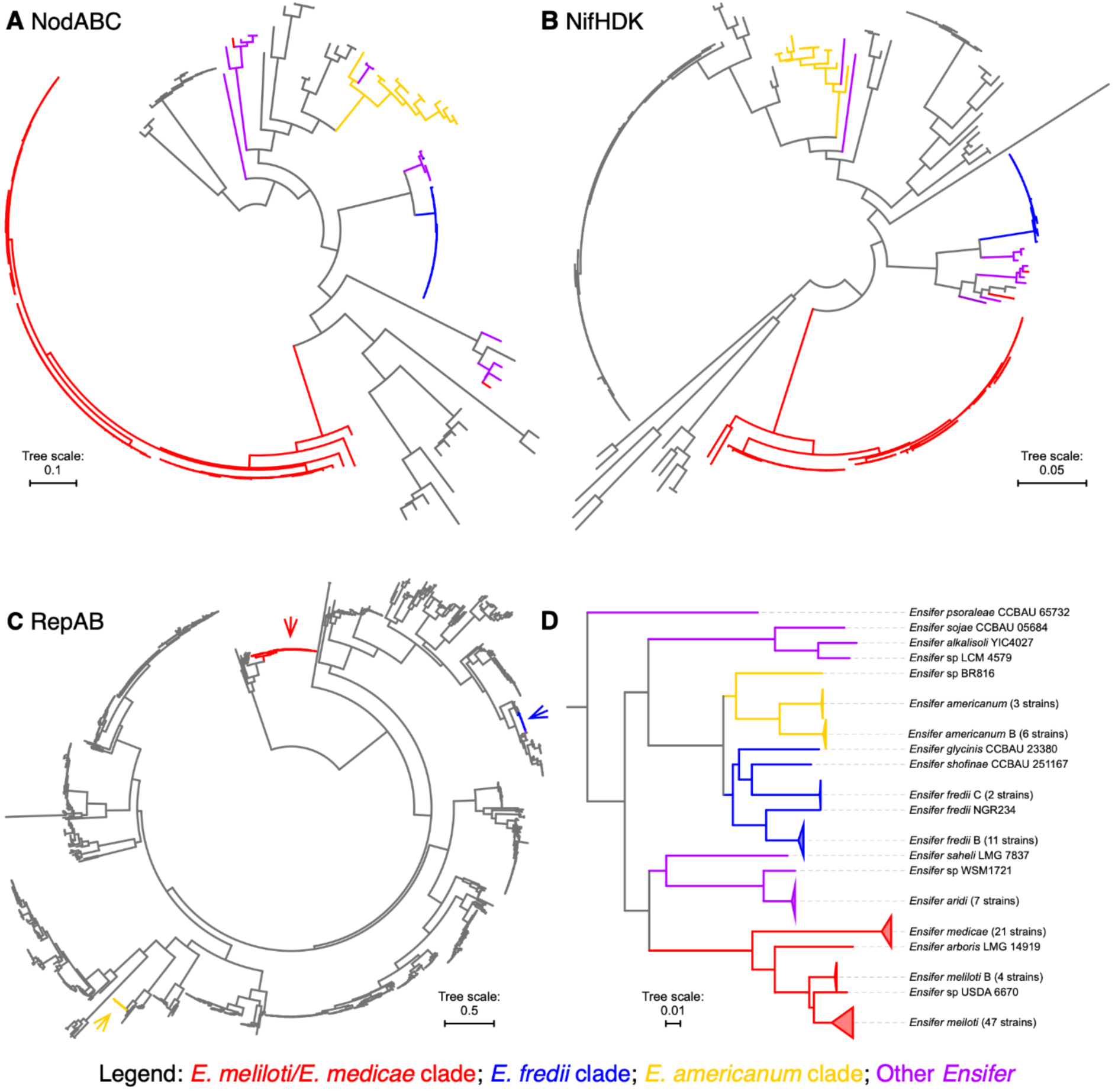
Evolution of SNF within the genus *Ensifer*. Maximum likelihood phylogenies of concatenated alignments of (**A**) NodABC nodulation proteins, (**B**) NifHDK nitrogenase proteins, and (**C**) RepAB replicon partitioning proteins of the order Rhizobiales. Branches corresponding to proteins from the genus *Ensifer* are indicated with colour. (**D**) A subtree of the whole-genome species phylogeny of Figure 1. Colours denote taxa whose symbiotic proteins are predicted to have been vertically acquired from a common ancestor. The scale bars represent the mean number of amino acid (A-C) or nucleotide (D) substitutions per site.

### The genomic features of the symbiotic and non-symbiotic clades differ

The pangenome of the 157 *Ensifer* strains was calculated to evaluate if there were global genomic differences between the symbiotic and non-symbiotic clades. Both clades had open pangenomes (Supplementary Figure S5). A PCA based on gene presence/absence revealed a clear separation of the two clades (Figure 3A), suggesting a divergence of the pangenomes of these clades. The symbiotic clade was sub-divided into two groups along the second component of the PCA (Figure 3A), which may suggest further levels of genomic separation. 2,130 genes were found in the core genomes of both clades, while 20% (542 genes) and 40% (1,377 genes) of the core genomes of the symbiotic and non-symbiotic clades, respectively, were absent from the core genome of the other clade; of these, about a third were completely absent from the other clade’s pangenome (Figure 3C). Of the 14,514 accessory genes (defined as genes found in at least 10% of at least one clade, excluding the 2,130 *Ensifer* core genes), only 2,352 (16%) were found in the pangenomes of both the symbiotic and non-symbiotic clades. This may suggest that these two clades predominately acquire new genes from distinct gene pools. Moreover, a statistically significant difference (Wilcoxon rank-sum test, *p* < 0.0001) in the genome sizes of the two clades was observed (Figure 3B); strains of the non-symbiotic clade carried 325 more genes, on average, than strains of the symbiotic clade (median difference of 470). Finally, based on the limited number of strains with finished genomes, strains of the symbiotic clade appear to generally have three copies of the rRNA operon whereas strains of the non-symbiotic clade appear to have a norm of five copies of their rRNA operon. Together, these multiple lines of data are consistent with there being a broad genomic divergence of the symbiotic and non-symbiotic clades of the genus *Ensifer*.

**Figure 3.**
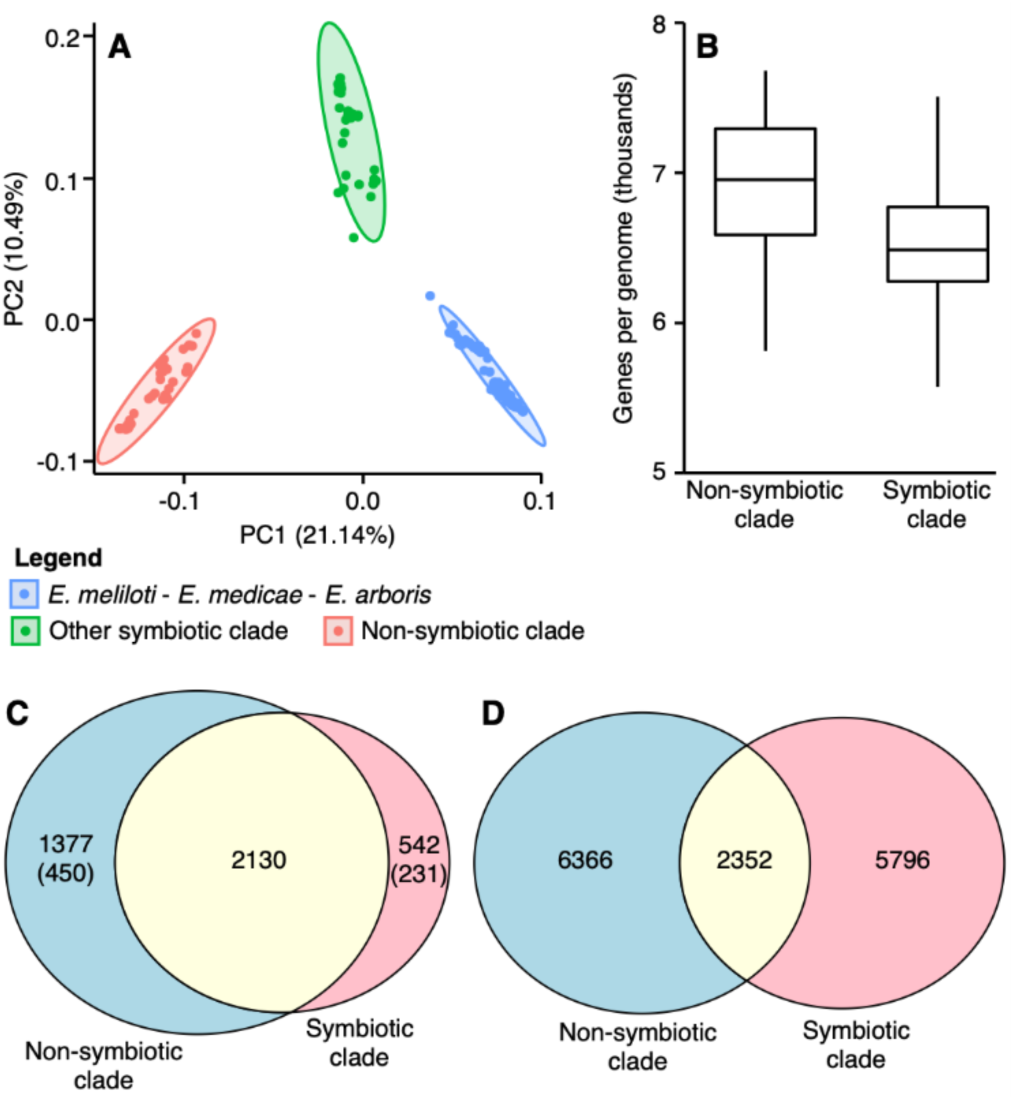
Global genome properties of the genus *Ensifer*. (**A**) A PCA plot based on the presence and absence of all orthologous protein groups in each of the 157 *Ensifer* strains. (**B**) Box-and-whisker plots displaying the number of genes per genome in the symbiotic and non-symbiotic *Ensifer* clades. (**C**) A Venn Diagram displaying the overlap in the core genomes of the symbiotic and non-symbiotic *Ensifer* clades. (**D**) A Venn Diagram displaying the overlap in the accessory genomes of the symbiotic and non-symbiotic *Ensifer* clades.

### Phenotypic features of the symbiotic and non-symbiotic clades differ

A subset of ten strains (Table 1), five each from the symbiotic and non-symbiotic clades, were subjected to a panel of assays to investigate how phenotypes vary across the genus *Ensifer*. No statistically significant differences were observed in the ability of members of the two clades to form biofilm or produce indol-3-acetic acid (Supplementary Figure S6, Supplementary Table S4). However, the two clades displayed a striking difference in their ability to grow in LB media; whereas strains of the non-symbiotic clade displayed robust growth in LB, strains of the symbiotic clade largely failed to grow (Figure 4A). Strains of the non-symbiotic clade also displayed a slightly faster specific growth rate, on average, than strains of the symbiotic clade in TY media (non-symbiotic clade: 0.54 ± 0.03 h^-1^; symbiotic clade: 0.44 ± 0.08 h^-1^; *p* = 0.03 from an ANOVA followed by Tukey’s test; Supplementary Figure S7A, Supplementary Table S5). On the other hand, strains of the symbiotic clade were, on average, better able to withstand heat stress (37°C) in TY media (Figure 4B). No statistically significant difference in the average ability of strains of the two clades to grow in minimal media with succinate or glucose as a carbon source, or with root exudates as a nitrogen source, was detected (Supplementary Figure S7, Supplementary Table S5).

**Table 1.**
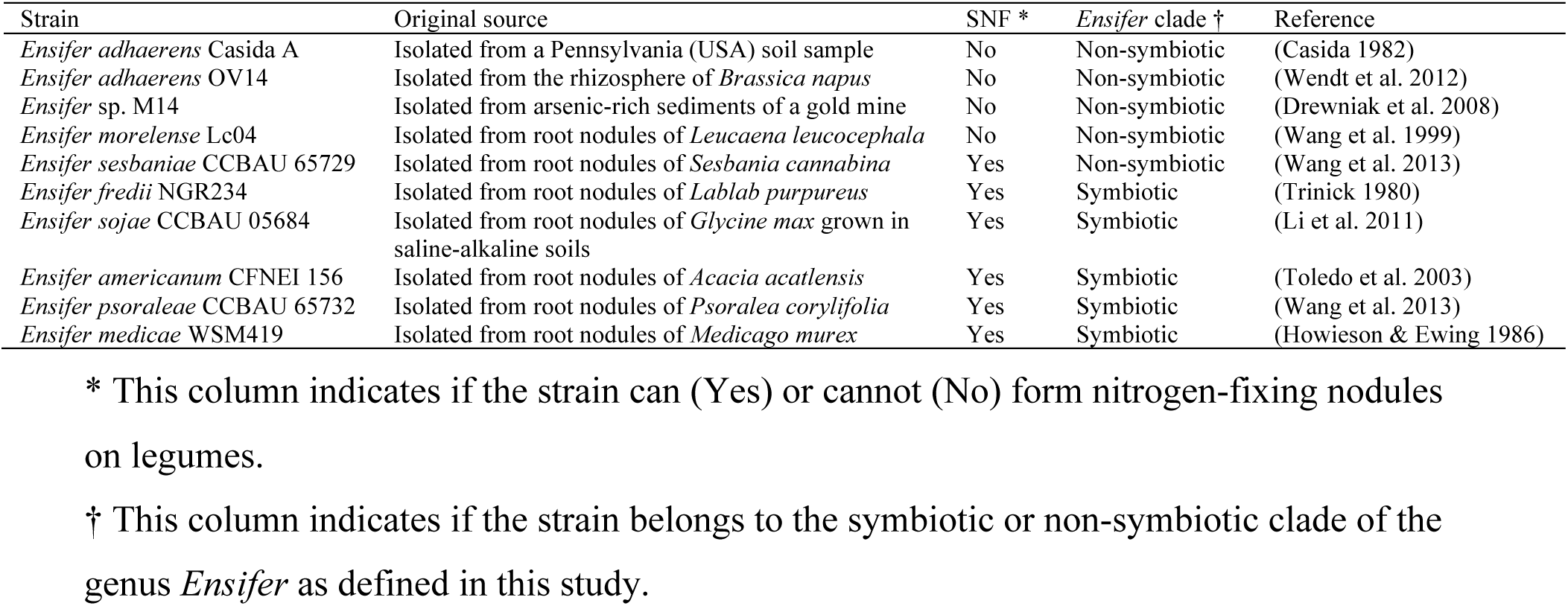
*Ensifer* strains phenotypically characterized in this study.

**Figure 4.**
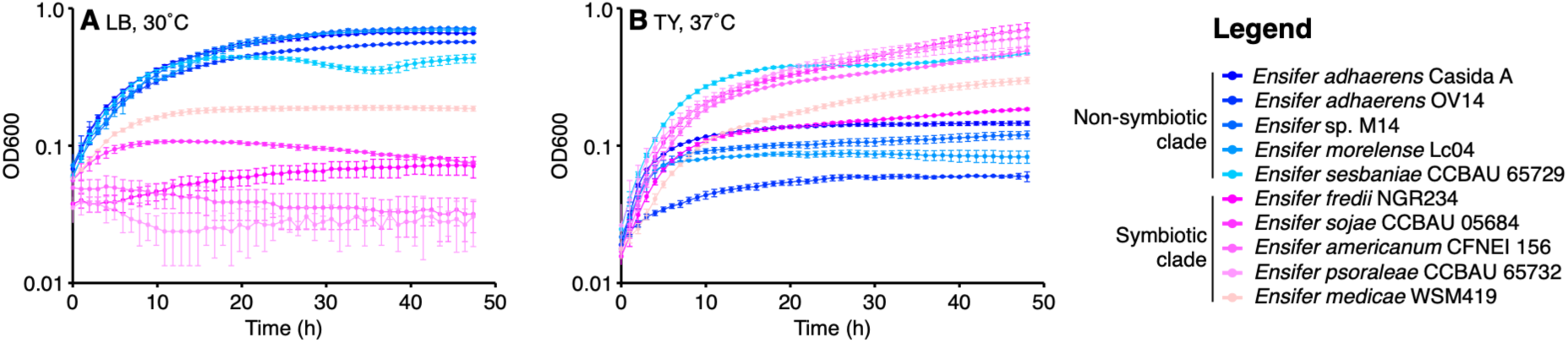
Growth properties of phylogenetically diverse *Ensifer* strains. *Ensifer* strains were grown in microplates without shaking. Data points represent the average of triplicate samples, while the error bars indicate the standard deviation. Shades of pink are used for strains of the symbiotic clade, while shades of blue are used for strains of the non-symbiotic clade. (**A**) Growth in LB medium at 30°C. (**B**) Growth in TY medium during heat stress (37°C).

The phenotypic properties of the genus *Ensifer* were further examined through evaluating the ability of the same ten strains to catabolize 190 carbon sources, and to grow in 96 osmolyte and 96 pH conditions, through the use of Biolog Phenotype MicroArrays™. Clustering the strains based on growth properties largely separated the symbiotic clade and non-symbiotic clade into distinct groups (Figure 5, Supplementary Figure S8). The exception was *E. sojae*, which formed its own intermediate group in the phenotype data. To aid in identifying which conditions best separate the symbiotic clade (including *E. sojae*) from the non-symbiotic clade, a linear discriminant analysis (LDA) was run over the AV values summarizing growth in each condition (Supplementary Dataset S4). In general, strains from the non-symbiotic clade better tolerated high pH (pH 9.0 to 9.5) than did strains from the symbiotic cluster. In contrast, strains of the symbiotic clade had better tolerance to low pH conditions (pH 3.5 to 4.5). In addition, the clades clearly differed in their overall metabolic abilities with strains of the non-symbiotic clade generally having a broader metabolic capacity than those of the symbiotic clade (Supplementary Table S6). Whereas strains of the symbiotic clade displayed robust growth on 65 carbon sources on average, strains of the non-symbiotic clade grew on an average of 81 carbon sources (*p* < 0.05, Student’s *t*-test). Overall, these experiments support the hypothesis that a variety of phenotypes, not just the ability to nodulate legumes, differ between the symbiotic and non-symbiotic clades of the genus *Ensifer*.

**Figure 5.**
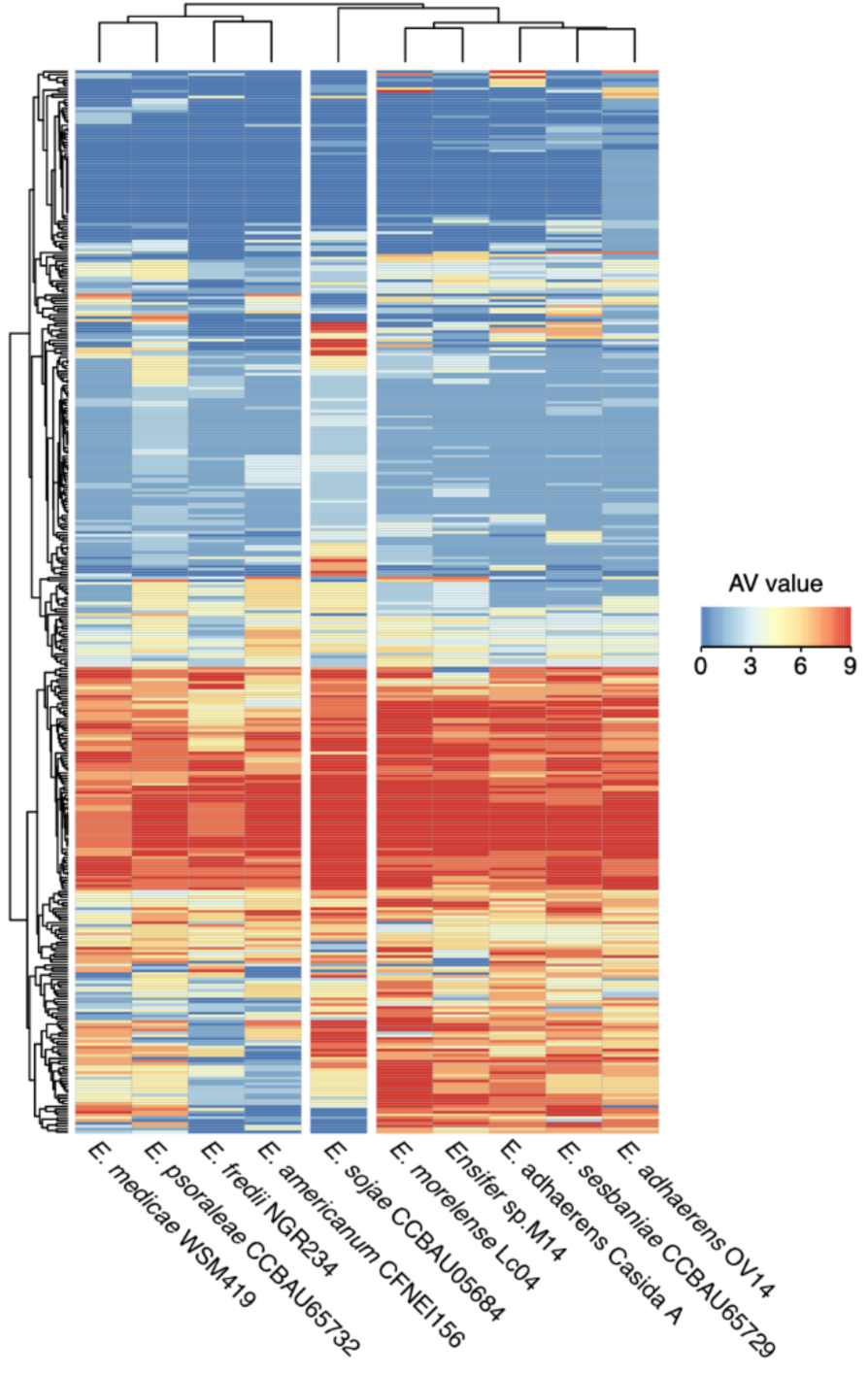
Phenotypic properties of phylogenetically diverse *Ensifer* strains. Ten *Ensifer* strains were screened for their ability to catabolize 190 carbon sources, and to grow in 96 osmolyte and 96 pH conditions using Biolog Phenotype MicroArray™plates PM1, PM2, PM9, and PM10. Growth in each well was summarized on a scale of 0 (dark blue) through 9 (dark red), with higher numbers representing more robust growth. A larger version of this figure, in which each condition is labelled along the Y-axis, is provided as Supplementary Figure S8.

## DISCUSSION

We investigated the evolution of SNF within the genus *Ensifer*, which includes a mix of nitrogen-fixing and non-nitrogen-fixing bacteria, as a model for the evolution of inter-kingdom mutualisms. Our results indicate that despite SNF having evolved multiple times within the genus *Ensifer*, the symbiotic and non-symbiotic strains are largely separated into two phylogenetic clades reminiscent of the general division of pathogenic and environmental strains between the genera *Burkholderia* and *Paraburkholderia* (Sawana et al. 2014). In addition to the prevalence of SNF, the two clades differed with respect to their genomic composition (pangenome content and genome size) and phenotypic properties (metabolic capacity and stress tolerance). There have been several revisions to the taxonomy of the genus *Ensifer*. Recently, a genome-based approach was proposed to standardize bacterial taxonomy (Parks et al. 2018) that splits the genus *Ensifer* into two genera: *Sinorhizobium* and *Ensifer*. The symbiotic and non-symbiotic clades identified here correspond with the genera *Sinorhizobium* and *Ensifer*, respectively, supporting the proposal to divide the genus *Ensifer* into two genera.

Our analyses revealed a complex evolutionary history of SNF within the genus *Ensifer*. In addition to SNF emerging a predicted six times or more, we detected possible losses of SNF and allele switches. Between one and six of the NodABC and NifHDK proteins were not detected in eight of the strains in the symbiotic clade (Figure 1). While this may be indicative of multiple losses of symbiosis, we cannot rule out that these are false negatives; most were draft genomes, and the one strain with a complete genome (*E. meliloti* M162) can nodulate 10 of 27 tested *Medicago truncatula* genotypes suggesting it does contain *nod* and *nif* genes (Sugawara et al. 2013). Based on the RepAB phylogeny (Figure 2), the symbiotic megaplasmid of *E. arboris* likely shares common ancestry with the symbiotic megaplasmids of the sister species *E. meliloti* and *E. medicae*. Yet, the nodulation and nitrogen fixation genes appeared distinct. Thus, we hypothesize that there was a recent replacement of the symbiotic genes in *E. arboris*, which is supported by the observation that *E. arboris* does not nodulate plants of the genus *Medicago* (Zhang et al. 1991), unlike *E. meliloti* and *E. medicae*.

The reason for the phylogenetic bias in the evolution of SNF within the genus *Ensifer* remains unclear, especially considering that strains from both clades are plant-associated (Bai et al. 2015). One possibility is that each clade occupies distinct niches within the soil and plant-associated environments. Indeed, the broader metabolic capacity of the non-symbiotic clade (Supplementary Table S6), which corresponded to a larger average genome size (Figure 3B), suggests these species are more capable of adapting to fluctuating nutritional environments. This is further supported by the apparently higher number of rRNA operons in strains of the non-symbiotic clade, which is generally thought to allow bacteria to more quickly respond to changing nutrient conditions (Stevenson & Schmidt 2004; Roller et al. 2016). Moreover, the non-symbiotic clade could be differentiated from the symbiotic clade based on its pangenome content (Figure 3A), which may suggest that strains of these clades acquire genes from distinct gene pools, further supporting the suggestion that they belong to distinct gene-cohesive groups and ecological niches. This hypothesis may then be interpreted in the framework of the stable ecotype model (*sensu* Cohan (Cohan 2006)), where the symbiotic and non-symbitic clades represent two, ecologically distinct and monophyletic groups and where periodic selection events (e.g. fitness for SNF) are recurrent.

An alternate, but not mutually exclusive, hypothesis is that the symbiotic *Ensifer* clade contains “facilitator” genes required to support SNF, similar to the theory that ancestral legumes contained a genetic “predisposition” necessary for the eventual evolution of rhizobium symbioses (Werner et al. 2014; Soltis et al. 1995; Doyle 2011). As we did not evaluate cause-and-effect relationships, our dataset does not definitely address this hypothesis. However, we observed numerous phenotypic and genotypic differences between the symbiotic and non-symbiotic clade, providing some support for this hypothesis. For example, the strains of the symbiotic clade appeared to have higher tolerance to low pH (Supplementary Figure S8, Supplementary Dataset S4), which is notable as the curled root hair is an acidic environment (Hawkins et al. 2017). At the genomic level, 231 of the core genes of the symbiotic clade were absent from the pangenome of the non-symbiotic clade and thus are good candidates as possible facilitators and follow-up studies. However, facilitators could also take the form of polymorphisms within highly conserved genes, as shown for *bacA* and the *Sinorhizobium* – *Medicago* symbiosis (diCenzo et al. 2017).

In summary, we show that the legume symbionts and non-symbionts of the genus *Ensifer* are largely segregated into two phylogenetically distinct clades that differ in their genomic and phenotypic properties. We suggest that these observations, which follow the guidelines recently reported for rhizobia and agrobacteria (de Lajudie et al. 2019), support the division of the genus into two genera: *Ensifer* for the non-symbiotic clade and *Sinorhizobium* for the symbiotic clade. However, formal descriptions and publication of the genera in the International Journal of Systematic and Evolutionary Microbiology (IJSEM) are still required. We also provide evidence that SNF evolved several times within this genus, but predominately within one monophyletic clade. These observations suggest that other genomic features (“facilitators”) aside from the core symbiotic genes could be required for the establishment of an effective symbiosis. These “facilitators” may reside in metabolic or genetic abilities present in the ancestral members of the symbiotic clades, which could have favoured the acquisition of effective SNF abilities by the members of this clade. This suggestion is supported by the ability to differentiate the strains of the two clades based on their pangenome content and phenotypic properties. However, as cause-and-effect relationships were not examined, follow-up study is required to more directly test this facilitators hypothesis.

## ACKNOWLEDGEMENTS

We are grateful to E. Martinez-Romero for providing strains *E. morelense* Lc04 and *E. morelense* Lc18, to E. Mullins (Teagasc, MTA2018233) for *E. adhaerens* OV14, to C.-F. Tian for *S. fredii* NGR 234, and to L. Dziewit for *Ensifer* sp. M14. AM was supported by the Fondazione Cassa di Risparmio di Firenze, grant n. 18204, 2017.0719, by the “MICRO4Legumes” grant (Italian Ministry of Agriculture), and by the grant “Dipartimento di Eccellenza 2018-2022” from the Italian Ministry of Education, University and Research (MIUR). LC was supported by the MICRO4Legumes grant (Italian Ministry of Agriculture). GCD was supported by a postdoctoral fellowship from the Natural Science and Engineering Research Council of Canada, and funding from Queen’s University.

