## Supplementary File S1 for "Symbiotic and non-symbiotic members of the genus *Ensifer* (syn. *Sinorhizobium*) are separated into two clades based on comparative genomics and high-throughput phenotyping"

Symbiotic and non-symbiotic members of the genus *Ensifer* (syn. *Sinorhizobium*) are separated into two clades based on comparative genomics and high-throughput phenotyping

Camilla Fagorzi[1], Alexandru Ilie[1], Francesca Decorosi[2], Lisa Cangioli[1], Carlo Viti[2], Alessio Mengoni[1*], George C diCenzo[1,3*]

[1] Department of Biology, University of Florence, Sesto Fiorentino, Italy

[2] Genexpress Laboratory, Department of Agriculture, Food, Environment and Forestry, University of Florence, Sesto Fiorentino, Italy

[3] Department of Biology, Queen's University, Kingston, Ontario, Canada

**Table S1.** Reference genomes used during scaffolding with MeDuSa.

| Genome assembly | Reference genomes | Assembly Accession | Citations |
|---|---|---|---|
| *E. psoraleae* CCBAU 65732 | *E. fredii* NGR234 | GCF_000018545.1 | (Schmeisser et al. 2009) |
| | *E. fredii* HH103 | GCF_002288485.1 | (Weidner et al. 2012) |
| | *E. fredii* CCBAU 83666 | GCF_000283895.1 | (Tian et al. 2012) |
| *E. sesbaniae* CCBAU 65729 | *E. adhaerens* OV14 | GCF_000583045.1 | (Rudder et al. 2014) |
| | *E. adhaerens* CasidaA | GCF_000697965.2 | (Williams et al. 2017) |
| *E. morelense* Lc04 | *E. adhaerens* OV14 | GCF_000583045.1 | (Rudder et al. 2014) |
| | *E. adhaerens* CasidaA | GCF_000697965.2 | (Williams et al. 2017) |
| *Rhizobium* sp. Lc18 | *R. tropici* CIAT 899 | GCF_000330885.1 | (Ormeño-Orrillo et al. 2012) |
| | *Rhizobium* sp. 10195 | GCF_002277895.1 | Unpublished |

**Table S2.** Hidden Markov models (HMMs) used for annotation of protein function.

| Protein annotation | Corresponding HMMs | Database |
|---|---|---|
| NodA | NodA | Pfam |
| | TIGR04245 | TIGRFAM |
| NodB | TIGR04243 | TIGRFAM |
| NodC | TIGR04242 | TIGRFAM |
| NifH | Fer4_NifH | Pfam |
| | TIGR01287 | TIGRFAM |
| NifD | TIGR01282 | TIGRFAM |
| | TIGR01860 | TIGRFAM |
| | TIGR01861 | TIGRFAM |
| NifK | TIGR01286 | TIGRFAM |
| | TIGR02931 | TIGRFAM |
| | TIGR02932 | TIGRFAM |
| RepA | TIGR03453 | TIGRFAM |
| RepB | RepB | Pfam |
| | TIGR00180 | TIGRFAM |
| | TIGR03454 | TIGRFAM |
| | TIGR03734 | TIGRFAM |

**Table S3.** Genome assembly summary statistics.

| | *E. psoraleae* CCBAU 65732 | *E. sesbaniae* CCBAU 65729 | *E. morelense* Lc04 | *Rhizobium* sp. Lc18 |
|---|---|---|---|---|
| Length | 7,427,611 | 6,897,201 | 7,073,540 | 7,125,406 |
| Scaffolds | 67 | 5 | 20 | 24 |
| G + C content | 61.23% | 62.09% | 61.75% | 59.91% |
| CDS | 7,001 | 6,457 | 6,544 | 6,656 |
| rRNA | 3 | 3 | 3 | 6 |
| tRNA | 56 | 55 | 52 | 48 |
| Average coverage | 89X | 103X | 75X | 78X |
| Scaffolds N50 | 3 | 1 | 2 | 2 |
| Scaffolds L50 | 1,137,413 | 3,711,528 | 1,461,079 | 1,454,936 |

**Table S4.** Statistical analysis of biofilm formation and IAA production by strains of the symbiotic and non-symbiotic clades of the genus *Ensifer*.

| Assay | Condition | Symbiotic clade * | Non-symbiotic clade † | *p*-value ¥ |
|---|---|---|---|---|
| Biofilm formation | TY, 30˚C | 0.15 ± 0.13 | 0.17 ± 0.07 | 0.71 |
| Biofilm formation | LB, 30˚C | 0.18 ± 0.25 | 0.07 ± 0.05 | 0.35 |
| IAA production | TSB, 30˚C | 3.76 ± 1.41 | 2.61 ± 1.79 | 0.29 |

* Strains of the symbiotic clade included in the assays were: *E. fredii* NGR234, *E. sojae* CCBAU 05684, *E. americanum* CFNEI 156, *E. psoraleae* CCBAU 65732, and *E. medicae* WSM419.

† Strains of the non-symbiotic clade included in the assays were: *E. adhaerens* Casida A, *E. adhaerens* OV14, *E.* sp. M14, *E. morelense* Lc04, and *E. sesbaniae* CCBAU 65729.

¥ The *p*-values were determined using ANOVA followed by Tukey's post-hoc tests.

**Table S5.** Statistical analysis of specific growth rates of strains of the symbiotic and non-symbiotic clades of the genus *Ensifer*.

| Medium | Temperature (˚C) | Symbiotic clade ($h^{-1}$) * | Non-symbiotic clade ($h^{-1}$) † | *p*-value ¥ |
|---|---|---|---|---|
| **TY** | **30** | **0.44 ± 0.08** | **0.54 ± 0.03** | **0.03** |
| TY | 37 | 0.38 ± 0.07 | 0.38 ± 0.13 | 1 |
| **LB** | **30** | **0.15 ± 0.20** | **0.43 ± 0.05** | **0.02** |
| LB | 37 | 0.38 ± 0.08 | 0.50 ± 0.16 | 0.18 |
| M9-succinate | 30 | 0.16 ± 0.06 | 0.21 ± 0.06 | 0.2 |
| M9-succinate | 37 | 0.24 ± 0.09 | 0.30 ± 0.10 | 0.39 |
| M9-glucose | 30 | 0.22 ± 0.04 | 0.23 ± 0.08 | 0.83 |
| M9-glucose | 37 | 0.25 ± 0.09 | 0.27 ± 0.10 | 0.73 |
| M9-succinate + root exudate ø | 30 | 0.15 ± 0.03 | 0.14 ± 0.07 | 0.63 |
| **Overall** | **-** | **0.26 ± 0.14** | **0.33 ± 0.16** | **0.03** |

* Strains of the symbiotic clade included in the assays were: *E. fredii* NGR234, *E. sojae* CCBAU 05684, *E. americanum* CFNEI 156, *E. psoraleae* CCBAU 65732, and *E. medicae* WSM419. Values represent the mean specific growth rate ± standard deviation.

† Strains of the non-symbiotic clade included in the assays were: *E. adhaerens* Casida A, *E. adhaerens* OV14, *E.* sp. M14, *E. morelense* Lc04, and *E. sesbaniae* CCBAU 65729. Values represent the mean specific growth rate ± standard deviation.

¥ The p-values were determined using ANOVA followed by Tukey's post-hoc tests.

ø Root exudate was provided as the sole source of nitrogen.

**Table S6.** Metabolic capacity of phylogenetically diverse *Ensifer* strains.

| Strain | Carbon sources supporting growth | |
|---|---|---|
| | AV ≥ 6 * | AV ≥ 4 * |
| Non-symbiotic clade | | |
| *E. adhaerens* Casida A | 80 | 106 |
| *E. adhaerens* OV14 | 69 | 86 |
| *Ensifer* sp. M14 | 86 | 98 |
| *E. morelense* Lc04 | 87 | 92 |
| *E. sesbaniae* CCBAU 65729 | 84 | 100 |
| **Average** | **81 ± 7** | **96 ± 8** |
| Symbiotic clade | | |
| *E. fredii* NGR234 | 56 | 70 |
| *E. sojae* CCBAU 05684 | 81 | 90 |
| *E. americanum* CFNEI 156 | 70 | 79 |
| *E. psoraleae* CCBAU 65732 | 50 | 71 |
| *E. medicae* WSM419 | 68 | 79 |
| **Average** | **65 ± 12** | **78 ± 8** |
| *p*-value † | ≤ 0.01 | 0.03 |

* AV values are a measure of the overall growth pattern of a strain in an individual well of a Biolog plate, measured on a scale of 0 (lowest growth) to 9 (best growth). Two thresholds were used to ensure that conclusions were not the result of choosing too strict or too lenient of a threshold. Numbers represent the number of wells (carbon sources) supporting growth at or above the indicated threshold; in the case of the averages, the mean ± the standard deviation is provided.

† The *p*-values are from Student's *t*-tests comparing the number of carbon sources supporting growth of the symbiotic and non-symbiotic clades.

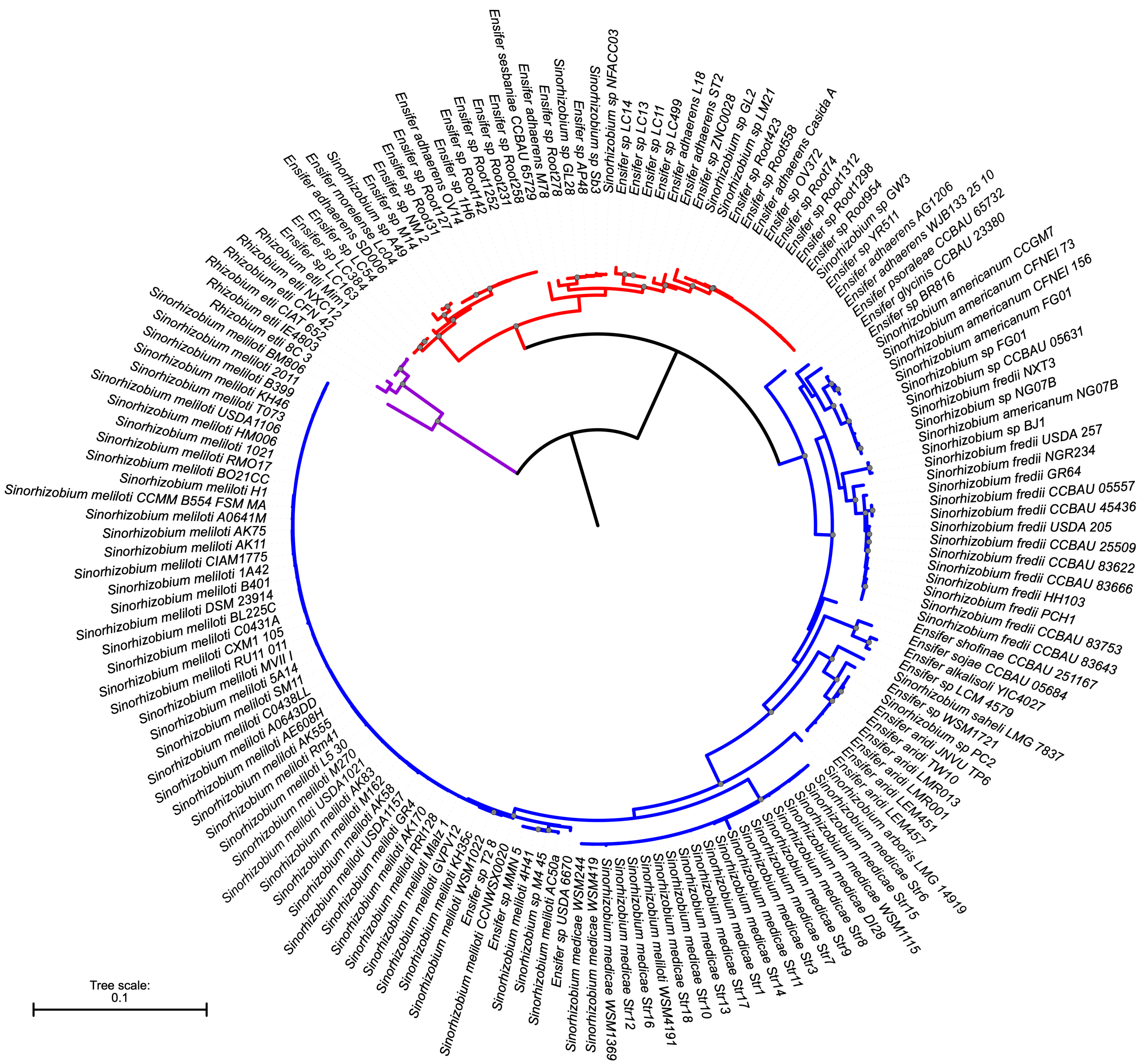


**Figure S1. Rooted phylogeny of the genus *Ensifer*.** A maximum likelihood phylogeny of 157 *Ensifer* strains, rooted with six *Rhizobium etli* strains, was prepared from a concatenated alignment of 30 highly conserved proteins (Frr, InfC, NusA, Pgk, PyrG, RplA, RplB, RplC, RplD, RplE, RplF, RplK, RplL, RplM, RplN, RplP, RplS, RplT, RpmA, RpoB, RpsB, RpsC, RpsE, RpsI, RpsJ, RpsK, RpsM, RpsS, SmpB, Tsf). Nodes with a bootstrap value of 90-100 are indicated with the gray dots. The scale represents the mean number of amino acid substitutions per site. The tree is colour-coded as follows: blue – the symbiotic clade; red – the non-symbiotic clade; purple – the *R. etli* outgroup. Strains are named as recorded in NCBI at the time of collection.

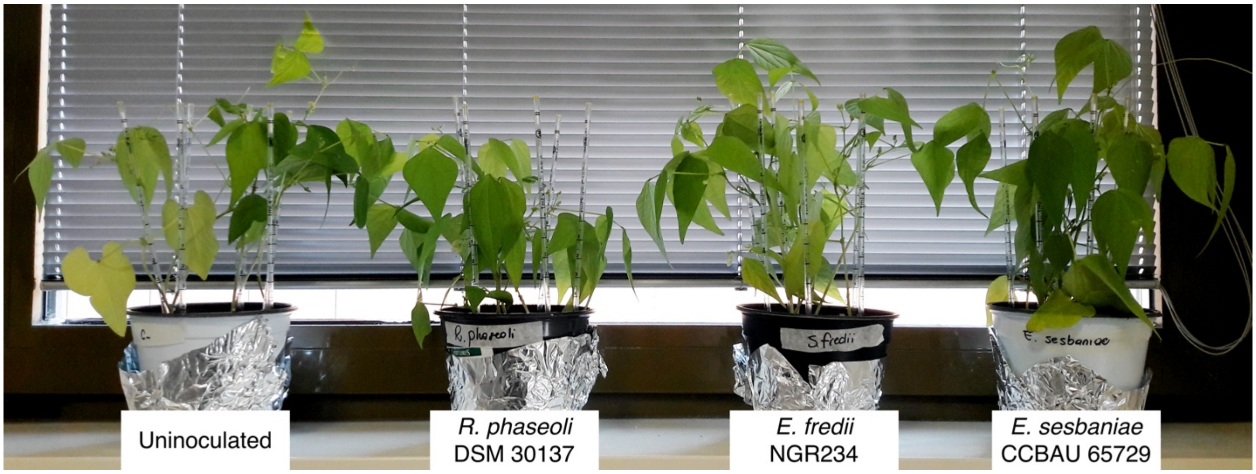


**Figure S2. Confirmation of the symbiotic abilities of *E. sesbaniae*.** *P. vulgaris* plants were either not inoculated or inoculated with *E. sesbaniae* CCBAU 65729, *E. fredii* NGR234, or *R. phaseoli* DSM30137. Pictures of the plants were taken four-weeks post-inoculation. Two to three small (2-3 mm) nodules per plant were detected on the inoculated plants but not on the uninoculated control. Unlike plants inoculated with any of the microsymbionts, uninoculated plants showed reduced growth and signs of chlorosis. Amplified Ribosomal DNA Restriction Analysis (ARDRA) confirmed that the nodules of all inoculated plants indeed contained the correct microsymbiont.

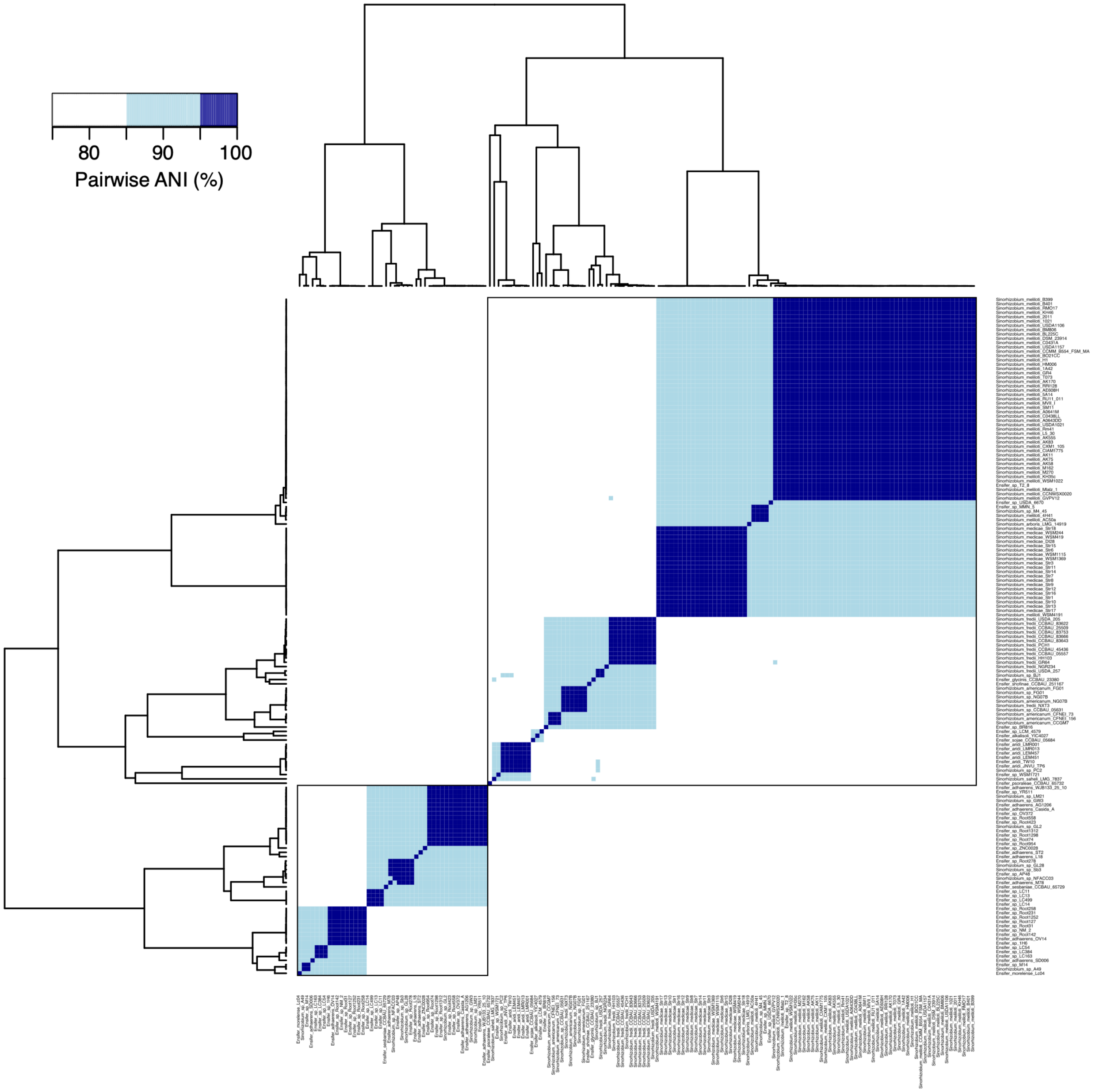


**Figure S3. ANI matrix of 157 *Ensifer* strains.** This figure displays the pairwise average nucleotide identity (ANI) values for each pair of genomes. Values were clustered along both axes using hierarchical clustering with average linkage and Pearson correlation distance. The large black box surrounds the symbiotic clade, while the smaller black box surrounds the non-symbiotic clade.

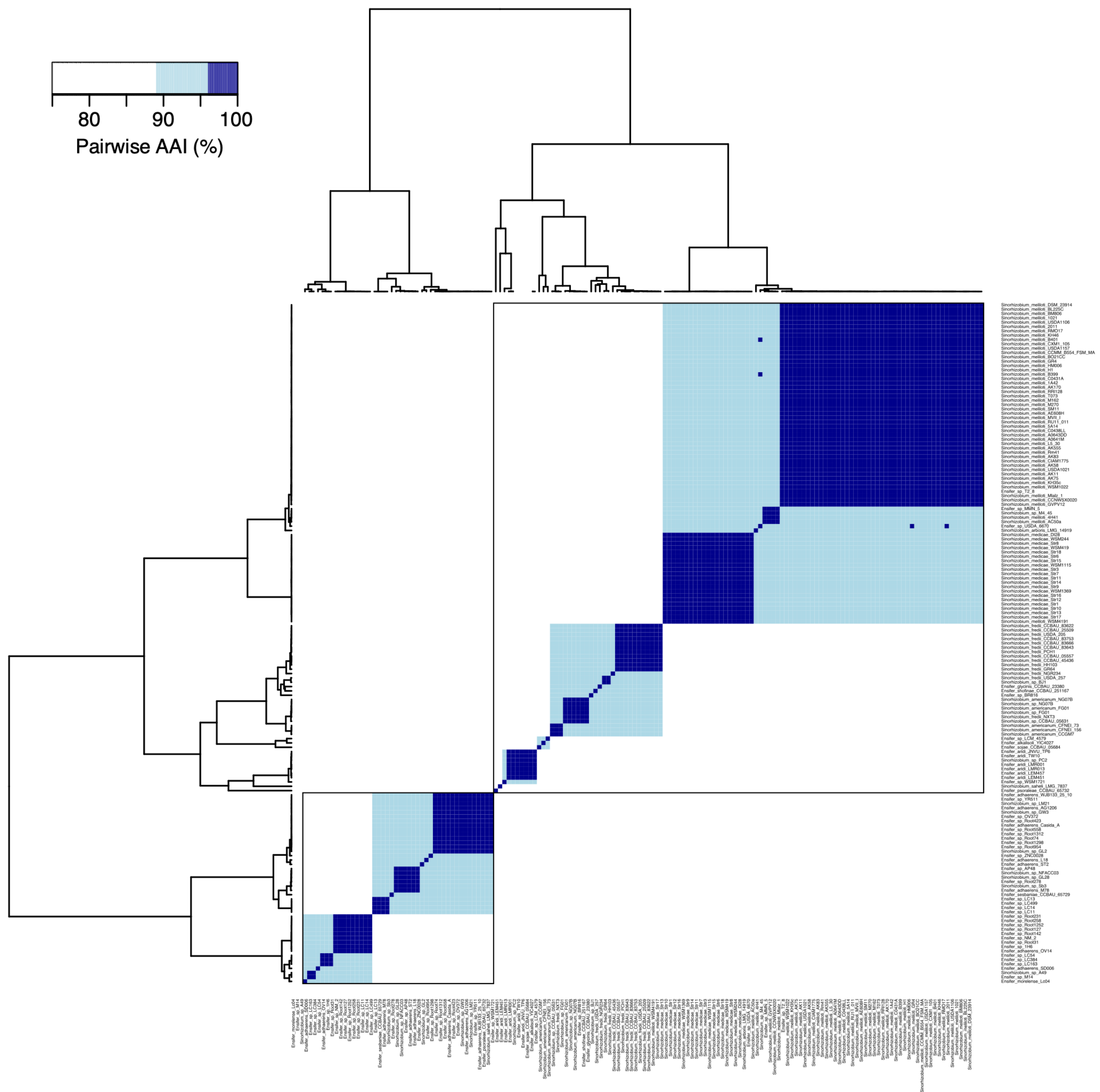


**Figure S4. AAI matrix of 157 *Ensifer* strains.** This figure displays the pairwise average amino acid identity (AAI) values for each pair of genomes. Values were clustered along both axes using hierarchical clustering with average linkage and Pearson correlation distance. The large black box surrounds the symbiotic clade, while the smaller black box surrounds the non-symbiotic clade.

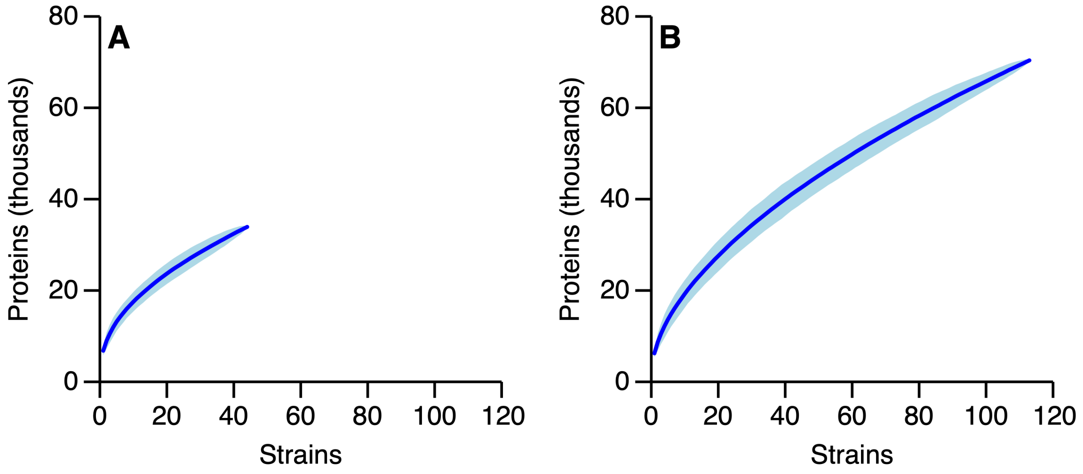


**Figure S5. Protein accumulation curves.** Protein accumulation curves are shown for the (**A**) non-symbiotic and (**B**) symbiotic clades of the genus *Ensifer*. Curves are based on 500 permutations.

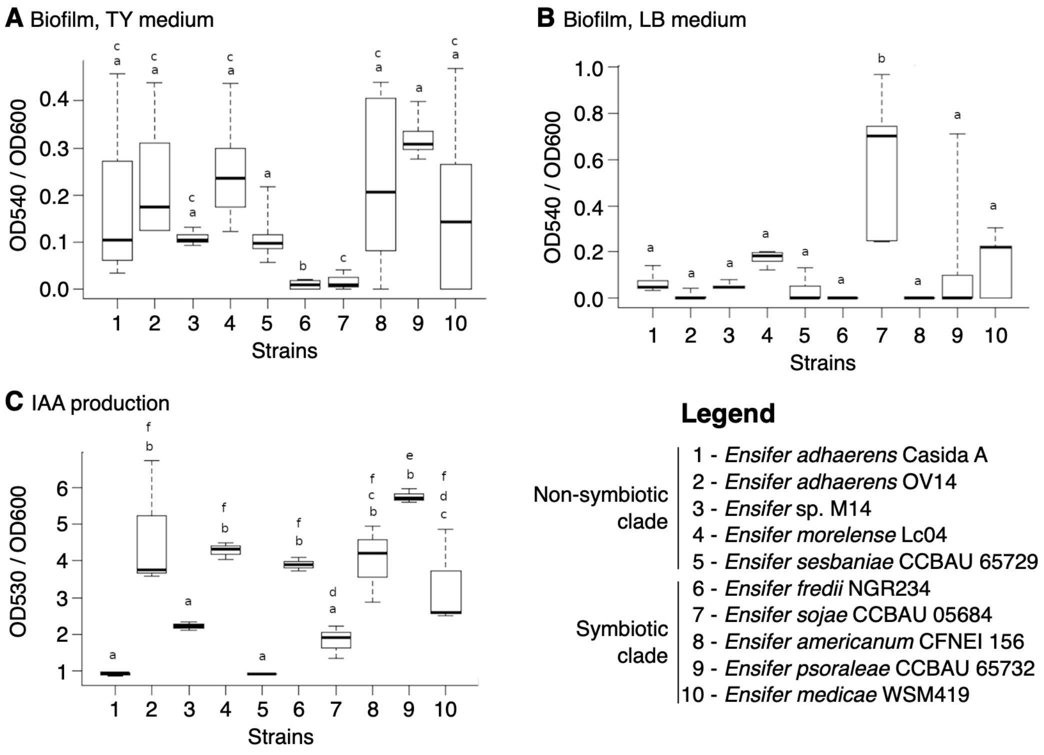


**Figure S6. Biofilm formation and IAA production by ten *Ensifer* strains.** All data are presented as box plots and represent the results from six (biofilm formation) or three (IAA production) replicates. The central think black lines indicate the median values. Letters indicate statistical groupings based on ANOVA followed by Tukey's post-hoc tests. Strain names corresponding to each number are given in the legend. (**A**) Biofilm production following growth in TY medium in microplates. (**B**) Biofilm production following growth in LB medium in microplates. (**C**) Indole-3-acetic acid (IAA) production following incubation in a 1:10 dilution of a TSB medium.

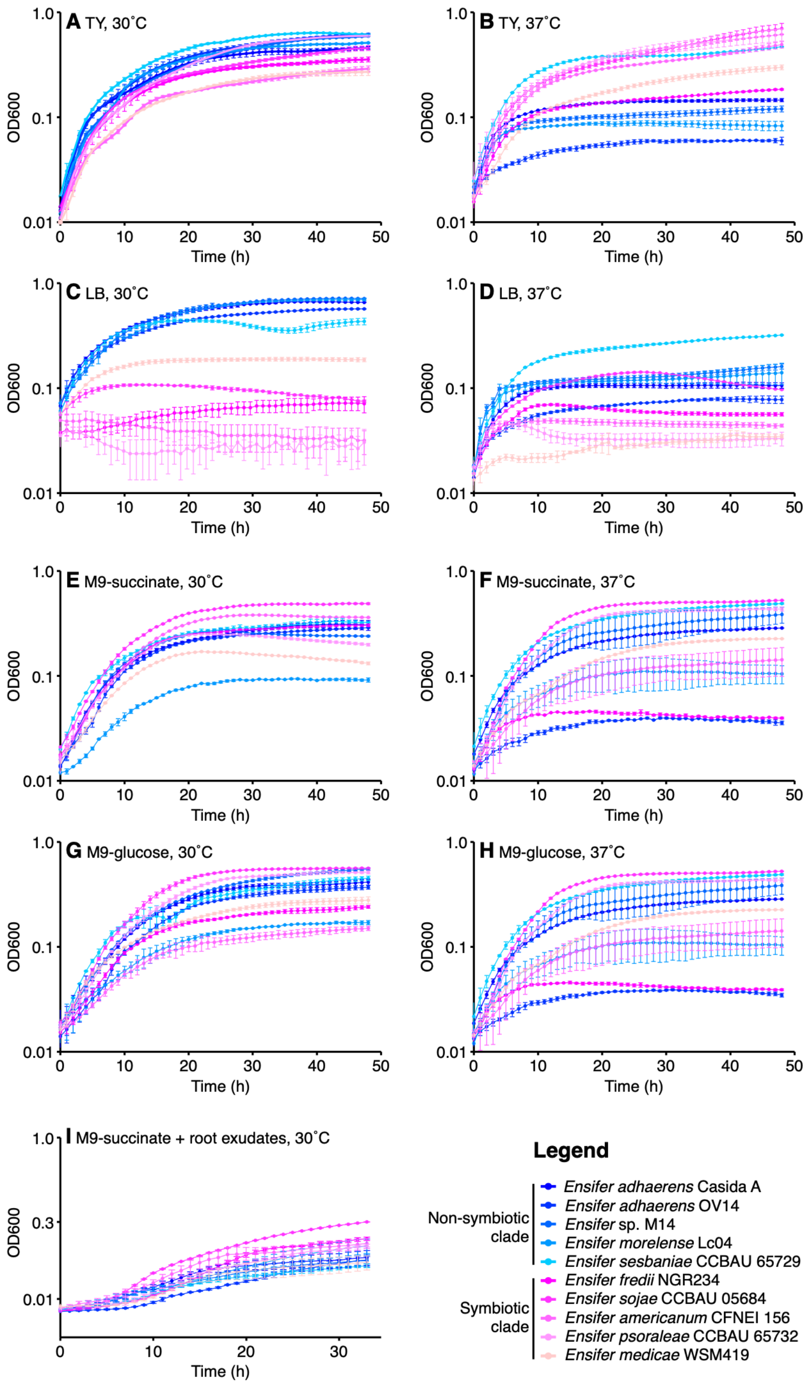


**Figure S7. Growth properties of ten *Ensifer* strains.** The *Ensifer* strains were grown in microplates without shaking. Data points represent the average of triplicate samples, while the error bars indicate the standard deviation. Shades of pink are used to represent strains from the symbiotic clade, while shades of blue are used to represent strains from the non-symbiotic clade. Strain names corresponding to each colour are given in the legend. (**A**) Growth in TY medium at 30˚C. (**B**) Growth in TY medium at 37˚C. (**C**) Growth in LB medium at 30˚C. (**D**) Growth in LB medium at 37˚C. (**E**) Growth in M9-succinate at 30˚C. (**F**) Growth in M9-succinate at 37˚C. (**G**) Growth in M9-glucose at 30˚C. (**H**) Growth in M9-glucose at 37˚C. (**I**) Growth in M9-succinate with root exudates as the sole source of nitrogen.

### REFERENCES (SUPPLEMENTARY MATERIALS)


Ormeño-Orrillo E et al. 2012. Genomic basis of broad host range and environmental adaptability of *Rhizobium tropici* CIAT 899 and *Rhizobium* sp. PRF 81 which are used in inoculants for common bean (*Phaseolus vulgaris* L.). BMC Genomics. 13:735.

Rudder S, Doohan F, Creevey CJ, Wendt T, Mullins E. 2014. Genome sequence of *Ensifer adhaerens* OV14 provides insights into its ability as a novel vector for the genetic transformation of plant genomes. BMC Genomics. 15:1–17.

Schmeisser C et al. 2009. *Rhizobium* sp. strain NGR234 possesses a remarkable number of secretion systems. Appl Environ Microbiol. 75:4035–4045.

Tian CF et al. 2012. Comparative genomics of rhizobia nodulating soybean suggests extensive recruitment of lineage-specific genes in adaptations. Proc Natl Acad Sci USA. 109:8629–8634.

Weidner S et al. 2012. Genome sequence of the soybean symbiont *Sinorhizobium fredii* HH103. J Bacteriol. 194:1617–1618.

Williams LE, Baltrus DA, O'Donnell SD, Skelly TJ, Martin MO. 2017. Complete genome sequence of the predatory bacterium *Ensifer adhaerens* Casida A. Genome Announc. 5:e01344–17.
