## Supplementary Figure S8 for "Symbiotic and non-symbiotic members of the genus *Ensifer* (syn. *Sinorhizobium*) are separated into two clades based on comparative genomics and high-throughput phenotyping"

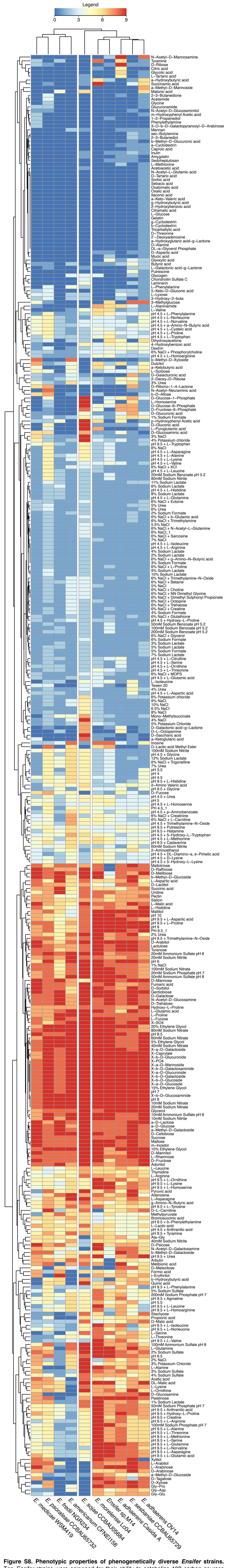


**Figure S8. Phenotypic properties of phenogenetically diverse *Ensifer* strains.** Ten *Ensifer* strains were screened for their ability to catabolize 190 carbon sources and to grow in 96 osmolyte and 96 pH conditions using Biolog Phenotype MicroArray plates PM1, PM2, PM9, and PM10. Overall growth in each well was summarized on a scale of 0 (dark blue) through 9 (dark red), with higher numbers representing more robust growth.
